# CoSTAR: Coarse Stem-Topology Alignment of Pseudoknotted RNA Structures by Relation-Constrained Search

**DOI:** 10.64898/2026.05.22.727263

**Authors:** Finn Archinuk, Hosna Jabbari

## Abstract

RNA structural alignment is a central task in comparative RNA analysis, but many efficient methods achieve tractability by restricting the class of admissible structures, often excluding pseudoknots. This exclusion is limiting for viral and regulatory RNAs, where conserved structure can remain informative even when sequence conservation is weak. We introduce a coarse RNA structural alignment algorithm that aligns secondary structures by searching over partial maps between stems rather than nucleotides. Each input structure is decomposed into stems, annotated with nucleotide-level features, and encoded by pairwise topological relations among stems. Alignment is formulated as a cost-minimizing partial stem map with skip operations, and the search tree is pruned by RNA-specific directionality and topological constraints derived from already aligned stems. For the stated cost function and over the class of injective, direction-preserving, topologically consistent stem maps, the search is exact. This shifts the dominant computational dependence from sequence length to the number and arrangement of stems. We evaluated the method on 2100 pairwise alignments sampled from seven Rfam families spanning 40–224 nucleotides and 2–15 stems. Across these benchmarks, the algorithm returned terminal coarse alignments in which every stem was either matched or skipped. We measured running time and search-tree width to characterize performance on diverse family-to-family comparisons. The experiments also show that ordering the input structures affects efficiency: using the structure with more stems as the search-driving structure reduces tree width. The resulting partial stem map is directly interpretable for RNA annotation and can be projected to nucleotide resolution for downstream sequence–structure analysis. The source code for CoSTAR is available at: https://github.com/TheCOBRALab/CoSTAR

## 1 Introduction

RNA molecules perform diverse informational, catalytic, and regulatory functions in the cell. Protein-coding mRNAs serve as templates for translation. Non-coding RNAs contribute to translation as ribosomal and transfer RNAs, to RNA processing as spliceosomal and small nucleolar RNAs, to gene regulation as small RNAs and long non-coding RNAs, to genome organization and defense, and to catalysis as ribozymes [19, 20, 4, 7]. Across these roles, RNA activity often depends not only on primary sequence, but also on structures formed within the molecule and through interactions with other RNAs, DNA, proteins, or small molecules [26, 5, 12]. Thus, the analysis of RNA function often requires methods that compare molecules beyond their primary sequences.

One important level of RNA structural organization is secondary structure, defined by the set of intramolecular base pairs in the molecule [26]. For many structural non-coding RNAs, functional constraints act on the base-pairing pattern at least as strongly as on the primary sequence. Consequently, homologous RNAs may show weak sequence identity while retaining corresponding base pairs. At a conserved base pair, a substitution at one paired position can be accompanied by a substitution at the other position so that base-pair complementarity is maintained. These coordinated changes are compensatory substitutions, and their statistical association across an alignment is referred to as covariation [10]. Comparative RNA methods use such covariation as evidence for conserved secondary structure [8]. Structural alignment is therefore a central computational primitive for comparing ncRNAs, annotating conserved structural elements, and transferring functional or mechanistic hypotheses between related RNAs [3].

The algorithmic difficulty of structural alignment depends strongly on the class of secondary structures handled. If all base pairs are non-crossing, the structure is pseudoknot-free and has a nested topology. This nesting property supports efficient dynamic programming decompositions, tree representations, and grammar-based models [24, 25, 9]. RNAforester, for example, uses a tree representation to compute local similarity between RNA secondary structures [15], whereas BEAR enriches dot-bracket strings with contextual structural symbols and learned substitution scores [18]. However, these efficient approaches are designed for a restricted structural class and do not preserve crossing base-pair dependencies. A pseudoknot occurs when two base pairs (*i, j*) and (*i*^′^, *j*^′^) satisfy *i < i*^′^ *< j < j*^′^ or *i*^′^ *< i < j*^′^ *< j*. Although pseudoknots constitute a minority of annotated structures in current datasets (bpRNA estimates that approximately 12% of annotated structures contain pseudoknots), they occur in important viral regulatory mechanisms, including gene expression and replication [6, 2]. Thus, the existing efficient structural comparison methods may miss the biologically important structural motifs that lie outside of their pseudoknot-free class of structures.

This limitation is especially relevant for viral non-coding and regulatory RNA elements. Viral genomes can accumulate substitutions rapidly; as sequence divergence increases, nucleotide-level alignments may fail to identify corresponding stems or structural motifs [21]. At the same time, particular stems and pseudoknotted motifs may be constrained by replication, translation, or regulatory function, and may therefore remain comparable even when the underlying nucleotides differ [2]. Some sequence–structure alignment methods can handle pseudoknotted inputs. LaRA 2, for example, aligns RNA sequences with pseudoknotted secondary structures [28]. Such methods return nucleotide-to-nucleotide alignments. In applications where the primary question is whether two RNAs contain corresponding stems, this resolution is not always necessary. We therefore consider a coarser alignment problem: compute a partial stem map between the stems of two RNA secondary structures, while preserving their relative topological relationships. This stem-level map can be used directly for structural annotation or as a set of anchors for subsequent nucleotide-level sequence–structure alignment.

Motivated by this setting, we formulate RNA structural alignment at the level of stems rather than nucleotides. For each input structure, short bulges and internal loops are first compressed according to a user-defined threshold, and the resulting structure is decomposed into stems. Each stem is represented by a feature vector that includes the number of base pairs, the number of nucleotides in its two stem arms, and its stacking-energy magnitude. We also precompute a relation array for each structure. For an ordered pair of stems (*A, B*), this array records the topological relation of *B* with respect to *A*: 5^′^-disjoint, 3^′^-disjoint, inside *A*, containing *A*, crossing *A* from the 5^′^ side, or crossing *A* from the 3^′^ side. Given two input structures *R*^*F*^ and *R*^*G*^, with ordered stem lists ℱ and 𝒢, a coarse alignment is a partial injective map from stems of ℱ to stems of 𝒢. Stems that are not included in the map are assigned skip costs, and matched stems are assigned costs derived from their feature vectors.

The computational bottleneck is the number of possible partial stem maps. If ℱ has *f* stems and 𝒢 has *g* stems, then, without any order constraint, the number of partial injective maps is 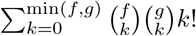. Indeed, for each *k*, one chooses *k* stems from ℱ, *k* stems from 𝒢, and an arbitrary bijection between the two chosen sets. CoSTAR restricts the search to direction-preserving maps: if *F*_*i*_ is matched before 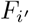, then its image in 𝒢 must also occur before the image of 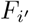. Under this restriction, the two chosen *k*-element subsets determine a unique order-preserving bijection. Hence the number of terminal direction-preserving partial maps, before applying topological constraints, is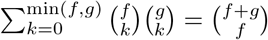, where the equality follows from Vandermonde’s identity. This space remains exponential in the number of stems, and the generated search tree also contains prefixes of such maps. CoSTAR therefore expands partial alignments in the 5^′^-to-3^′^ order of ℱ. When assigning the next stem *F*_*i*_, a candidate *G*_*j*_ is considered only if *j* is larger than every previously matched index in 𝒢. The candidate is admissible only if, for every previously matched pair (*F*_*p*_, *G*_*q*_), *ρ*_*F*_ (*p, i*) = *ρ*_*G*_(*q, j*), where *ρ*_*F*_ (*p, i*) and *ρ*_*G*_(*q, j*) denote the corresponding ordered stem relations in the two input structures. These precomputed local consistency checks reduce the branching factor without requiring either structure to be pseudoknot-free.

### Contributions

We introduce CoSTAR, a relation-constrained search algorithm for coarse stem-topology alignment of RNA secondary structures that may contain pseudoknots. CoSTAR represents each input as an ordered list of stems with nucleotide-derived features and precomputed pairwise topological relations. It then solves a minimum-cost alignment problem over partial injective stem maps with explicit skip operations, using 5^′^-to-3^′^ order and relation-array consistency to prune infeasible extensions. We evaluate CoSTAR on 2100 pair-wise alignments from seven Rfam families with 2–15 stems, characterizing running time and search-tree width and showing that placing the structure with more stems in the search-driving role empirically reduces search-tree width. The output is an interpretable partial stem map for annotation or downstream nucleotide-level sequence–structure alignment.

## 2 Related Work

RNA comparison methods differ in the information they use, the structural classes they support, and the resolution of the output. Classical sequence–structure alignment methods extend sequence alignment by incorporating base-pairing information into the objective. The Sankoff formulation gives a general framework for simultaneous alignment and folding, but its full dynamic program is computationally expensive [24]. LocARNA implements an efficient Sankoff-style approach for local sequence–structure alignment of pseudoknot-free RNAs and has been used for structure-based clustering of non-coding RNAs [27]. LaRA and LaRA 2 address fixed sequence–structure alignment using combinatorial optimization. In particular, LaRA models sequence–structure alignment as an integer linear program, and LaRA 2 accelerates pairwise alignment by using parallel execution and vectorized alignment kernels [1, 28]. These methods are appropriate when the desired output is a nucleotide-to-nucleotide alignment. Our objective is different: given two fixed structures, we compute a partial stem map between stems. This coarser output records which stems correspond and which are skipped, and can later be projected to nucleotide resolution if needed.

Several methods compare RNA secondary structures primarily by transforming or enriching structural representations. RNAforester represents pseudoknot-free secondary structures as ordered forests and computes local or global structural alignments [15]. BEAR replaces standard dot-bracket notation with a context-aware alphabet that distinguishes paired and unpaired positions according to their structural context, and then compares the resulting strings using learned substitution scores [18]. More recently, bpRNA-align uses feature-specific structural arrays, context-dependent affine gap penalties, and a structural substitution matrix to compute global similarity scores between RNA secondary structures [16]. These approaches provide useful structural similarity measures, but their alignments are defined over nucleotide positions or position-level structural symbols. In contrast, our method searches directly over partial maps between stems, which are the objects returned by the output alignment.

Pseudoknotted structures require representations that do not rely on purely nested topology. ASPRAlign compares secondary structures with arbitrary pseudoknots by transforming them into algebraic or structural RNA trees and computing the ASPRA distance [22]. This provides an efficient structure-only distance for pseudoknotted RNAs. However, the ASPRA distance is a global dissimilarity measure rather than an explicit feature-weighted correspondence between stems. Our method instead keeps pseudoknotted relations as pairwise topological constraints during search, while retaining stem-level quantitative features such as stem length and stacking energy.

Descriptor-based RNA motif search provides another relevant abstraction. In RNArobo, a user specifies a motif descriptor containing sequence and structural constraints, and the algorithm searches genomic sequences for matching regions [23]. Although descriptor search is not a pairwise alignment problem, it illustrates the utility of representing RNA structure in terms of ordered helices and other structural descriptors rather than individual nucleotides. Our reduced topology graph follows this element-level view, but uses it to align two complete structures rather than to search a sequence database for occurrences of a query descriptor.

Topology-based abstractions are also closely related. RNA abstract shapes collapse secondary structures into branching forms, enabling clustering of structures that differ in local base-pair details but share a global organization [13]. RNA-as-Graphs represents RNA secondary structures as graph topologies, including pseudoknotted structures through dual graphs [11]. Such abstractions are intentionally non-injective: distinct structures can map to the same abstract object when they differ in stem length, nucleotide composition, or energetic stability but not in topology. Super N-motifs takes a different alignment-free approach by decomposing structures into simple motifs and constructing vector representations from informative motif combinations [14]. These representations are useful for clustering or large-scale comparison, but they do not directly return a one-to-one correspondence between stems in two RNAs.

Our method occupies an intermediate point between nucleotide-level alignment and alignment-free topology comparison. It supports pseudoknotted inputs, uses topology to constrain the search, and uses nucleotide-derived stem features to avoid purely topological degeneracy. The resulting output is an explicit partial map between stems, rather than only a nucleotide alignment, a string alignment, or a scalar distance. This partial map can be projected to a nucleotide-level alignment which we will use to demonstrate alignment quality.

## 3 Data Representation

In this section we define the representation used by our CoSTAR algorithm. The input is an RNA sequence together with a fixed secondary structure. The output of the preprocessing step is an ordered stem list, a reduced topology graph, a relation array over ordered pairs of stems, and a feature vector for each stem.

### 3.1 RNA sequence and secondary structure

An RNA sequence *S* = *S*_1_, …, *S*_*n*_ is a string over the alphabet {*A, C, G, U* }. The indices 1, …, *n* are ordered from the 5^′^ end to the 3^′^ end. For 1 ≤ *i* ≤ *j* ≤ *n*, let *S*_*i,j*_ = *S*_*i*_, …, *S*_*j*_ denote the subsequence of *S*, and let [*i, j*] denote the corresponding interval of nucleotide positions.

A base pair of *S* is an ordered pair (*i, j*), where 1 ≤ *i < j* ≤ *n*. In a canonical sequence-compatible structure, *S*_*i*_ and *S*_*j*_ are complementary, i.e., {*S*_*i*_, *S*_*j*_} is one of {*A, U* }, {*C, G*}, or {*G, U* }. The alignment algorithm below is specified for the base-pair set itself, so annotated non-canonical pairs can also be admitted provided that the pairing relation is well defined. A secondary structure *R* for *S* is a set of base pairs such that each nucleotide participates in at most one pair:

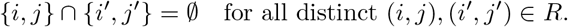

A secondary structure is pseudoknot-free if it does not contain two base pairs (*i, j*) and (*i*^′^, *j*^′^) such that *i < i*^′^ *< j < j*^′^ or *i*^′^ *< i < j*^′^ *< j*. Otherwise, the structure is pseudoknotted. Pseudoknotted structures are admissible inputs to our method.

### 3.2 Stems and bulge compression

A stem is a maximal chain of base pairs that are consecutive, or nearly consecutive after ignoring a bounded number of unpaired nucleotides. Let *β* ≥ 0 be a user-defined bulge compression parameter. Consider two base pairs (*i, j*) and (*k, ℓ*) in *R*, with *i < k < ℓ < j*. They are stacked if *k* = *i*+1 and *ℓ* = *j* − 1. They are *β*-adjacent if all nucleotides in the intervals [*i* + 1, *k* − 1] and [*ℓ* + 1, *j* − 1] are unpaired in *R*, and (*k* − *i* − 1) + (*j* − *ℓ* − 1) ≤ *β*. Thus, *β* = 0 recovers the ordinary definition of consecutive stacked base pairs. Larger values of *β* allow short bulges or internal loops to be compressed when defining coarse stems. The original sequence and base-pair set are not changed; the parameter *β* only determines which base pairs are grouped into the same stem. As default we set *β* = 2.

A stem *h* is a maximal ordered sequence of base pairs *h* = (*i*_1_, *j*_1_), (*i*_2_, *j*_2_), …, (*i*_*r*_, *j*_*r*_) such that *i*_1_ *< i*_2_ *<* …*< i*_*r*_ *< j*_*r*_ *<* …*< j*_2_ *< j*_1_, and (*i*_*t*−1_, *j*_*t*−1_) and (*i*_*t*_, *j*_*t*_) are *β*-adjacent for every 1 ≤ *t* ≤ *r*. The integer *r* is the number of base pairs in the stem.

Each stem has two arms. The 5^′^ arm of *h* is the interval *h*^−^ = [*i*_1_, *i*_*r*_], and the 3^′^ arm is the interval *h*^+^ = [*j*_*r*_, *j*_1_]. We refer to the ordered pair (*h*^−^, *h*^+^) as the arm representation of *h*. The 5^′^ arm always precedes the 3^′^ arm in the sequence order.

Let ℋ (*R, β*) = (*h*_1_, …, *h*_*M*_) be the ordered stem list obtained from *R* using compression parameter *β*, where stems are ordered by the left endpoint of their 5^′^ arms, and *M* is the total number of stems. When *β* is fixed by context, we write simply ℋ (*R*).

### 3.3 Reduced topology graph

The reduced topology graph records the order of stem arms along the RNA backbone and the pairing relationship between the two arms of each stem. Let 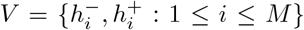 be the set of arm nodes. Sort these 2*M* intervals by their left endpoint in the 5^′^-to-3^′^ order, and denote the resulting sequence by *v*_1_, *v*_2_, …, *v*_2*M*_ . The reduced topology graph is *T*_*R*_ = (*V, E*_bb_ ∪ *E*_stem_), where *E*_bb_ = {(*v*_*t*_, *v*_*t*+1_) : 1 ≤ *t <* 2*M* } is the set of directed backbone edges, and 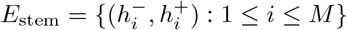 is the set of stem edges. Unpaired regions that do not belong to compressed stems are not represented as vertices. Hence *T*_*R*_ has 2*M* vertices, 2*M* − 1 backbone edges, and *M* stem edges. This representation is related to descriptor-based RNA motif representations, where structures are described through ordered helices and intervening regions [23]; here we retain only the stem arms and their order because the alignment algorithm operates on stems rather than on unstructured intervals. The graph *T*_*R*_ is used to define and visualize the reduced topology; the search algorithm uses the ordered stem list, the feature vectors, and the relation array induced by *T*_*R*_.

### 3.4 Pairwise topological relations

The alignment algorithm uses pairwise topological relations between stems as constraints. We first define an ordering relation on arm intervals. For two arm intervals *x* and *y*, write *x* ≺ *y* if interval *x* ends before interval *y* begins.

Let *A* = (*a, a*^′^) and *B* = (*b, b*^′^) be two distinct stems in arm representation, where *a* and *b* are the 5^′^ arms and *a*^′^ and *b*^′^ are the 3^′^ arms. We use the ordered relation-label tuple Λ = (*λ*_1_, *λ*_2_, *λ*_3_, *λ*_4_, *λ*_5_, *λ*_6_) defined by the following mutually exclusive cases:

| $\ell$ | condition | relation of $B$ with respect to $A$ | code alias |
| --- | --- | --- | --- |
| 1 | $b' \prec a$ | 5'-disjoint | 5'SL |
| 2 | $b \prec a \prec b' \prec a'$ | 5'-crossing | 5'PK |
| 3 | $b \prec a \prec a' \prec b'$ | contains | outer |
| 4 | $a \prec b \prec a' \prec b'$ | 3'-crossing | 3'PK |
| 5 | $a' \prec b$ | 3'-disjoint | 3'SL |
| 6 | $a \prec b \prec b' \prec a'$ | inside | inner. |

The semantic labels are used in the text. The code aliases are retained only for consistency with the implementation and figures.

For a structure *R* with ordered stem list ℋ(*R*) = (*h*_1_, …, *h*_*M*_), define *ρ*_*R*_(*i, j*) as the unique label in Λ giving the relation of stem *h*_*j*_ with respect to stem *h*_*i*_, for *i* ≠ *j*. For any ordered pair of distinct stems, exactly one label in Λ applies. The reciprocal relation is obtained by reversing the ordered pair. For example, if *ρ*_*R*_(*i, j*) = *λ*_1_, then *ρ*_*R*_(*j, i*) = *λ*_5_; if *ρ*_*R*_(*i, j*) = *λ*_3_, then *ρ*_*R*_(*j, i*) = *λ*_6_; and if *ρ*_*R*_(*i, j*) = *λ*_2_, then *ρ*_*R*_(*j, i*) = *λ*_4_.

We precompute *ρ*_*R*_ for all ordered pairs of stems. The relation array for structure *R* is the Boolean tensor Γ_*R*_ ∈ {0, 1}^*M*×*M*×6^, where for *i, j* ∈ {1, …, *M* } and *ℓ* ∈ {1, …, 6},

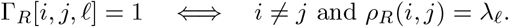

Diagonal entries Γ_*R*_[*i, i, ℓ*] are set to 0 and are not used. Since there are *M* (*M* −1) ordered pairs of distinct stems, the relation array is computed in *O*(*M* ^2^) time and stored in *O*(*M* ^2^) space. This array is independent of the second structure being aligned, so it can be cached and reused across multiple alignments involving the same RNA structure.

### 3.5 Stem features

The reduced topology graph and relation array encode the relative arrangement of stems, but they do not distinguish stems with the same topological relation profile and different physical properties. Therefore each stem is also assigned a feature vector derived from the original nucleotide-level structure.

For a stem *h* = (*i*_1_, *j*_1_), …, (*i*_*r*_, *j*_*r*_), we define three features. The first is the number of base pairs, *ϕ*_1_(*h*) = *b*. The second is the number of nucleotide positions between the two stem arms, *ϕ*_2_(*h*) = *j*_*r*_ − *i*_*r*_ − 1. This value provides geometric spacing considerations. The third is the magnitude of the stacking free energy contributed by adjacent stacked base-pair steps in the stem. Let *e*_*s*_(*i, j*) denote the nearest-neighbor stacking free energy for the adjacent stacked base-pair step (*i, j*), (*i* + 1, *j* − 1), using standard thermodynamic parameters [17]. Then

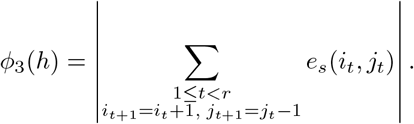

If no adjacent stacked base-pair step occurs in *h*, then *ϕ*_3_(*h*) = 0. The absolute value is used so that larger feature values correspond to larger stabilizing contributions.

The feature vector of *h* is *ϕ*(*h*) = (*ϕ*_1_(*h*), *ϕ*_2_(*h*), *ϕ*_3_(*h*)) . All feature vectors are computed once during preprocessing and are used later to define match and skip costs in the alignment algorithm.

### 3.6 Preprocessing output

For an input sequence–structure pair (*S, R*) and bulge compression parameter *β*, the preprocessing step returns

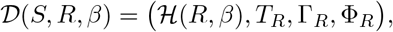

where ℋ(*R, β*) is the ordered stem list, *T*_*R*_ is the reduced topology graph, Γ_*R*_ is the relation array, and Φ_*R*_ = (*ϕ*(*h*_1_), …, *ϕ*(*h*_*M*_)) is the list of stem feature vectors. The alignment algorithm in the next section takes two such preprocessed representations as input.

## 4 CoSTAR Algorithm

For the stated cost function and over the class of injective, direction-preserving, topologically consistent stem maps, the CoSTAR search is exact. We now define the coarse alignment problem and the CoSTAR algorithm. Let (*S*^*F*^, *R*^*F*^) and (*S*^*G*^, *R*^*G*^) be two input sequence–structure pairs, and let

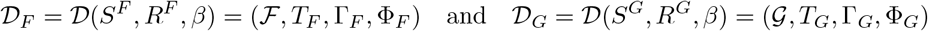

be their preprocessed representations. Here ℱ = (*F*_1_, …, *F*_*f*_) and 𝒢 = (*G*_1_, …, *G*_*g*_) are the ordered stem lists of the two structures.

### 4.1 Coarse alignments

A coarse alignment is a partial one-to-one correspondence between stems of ℱ and stems of 𝒢. Formally, let *I*_*F*_ = {1, …, *f* } and *I*_*G*_ = {1, …, *g*} be the stem-index sets of the two structures. A partial map

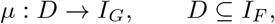

is injective if *µ*(*i*) ≠ *µ*(*i*^′^) for all distinct *i, i*^′^ ∈ *D*. We interpret *µ*(*i*) = *j* as matching stem *F*_*i*_ to stem *G*_*j*_. Stems in *I*_*F*_ \ *D* and *I*_*G*_ \ *µ*(*D*) are skipped.

The search is restricted to direction-preserving maps. A partial map *µ* is direction-preserving if, for all *i, i*^′^ ∈ *D, i < i*^′^ ⇒ *µ*(*i*) *< µ*(*i*^′^). This constraint preserves the 5^′^-to-3^′^ order of matched stems. It does not require the structures to be pseudoknot-free; pseudoknotted relationships are represented separately by the pairwise relation arrays.

A direction-preserving map is topologically consistent if every pair of matched stems has the same ordered relation in both structures. A partial map *µ* is topologically consistent if *ρ*_*F*_ (*p, i*) = *ρ*_*G*_(*µ*(*p*), *µ*(*i*)) for every distinct *p, i* ∈ *D*. Equivalently, Γ_*F*_ [*p, i*, ·] = Γ_*G*_[*µ*(*p*), *µ*(*i*), ·] for every ordered pair of matched stems. The alignment problem is to find a minimum-cost partial map that is injective, direction-preserving, and topologically consistent.

### 4.2 Match and skip costs

The cost of matching *F*_*i*_ to *G*_*j*_ is the weighted *ℓ*_1_ distance between their feature vectors:

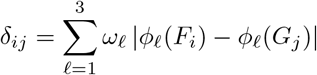

where *ω*_1_, …, *ω*_*d*_ ≥ 0 are feature weights (with default values of 1). All pairwise match costs are precomputed in an *f* × *g* matrix Δ = (*δ*_*ij*_).

Skipping a stem is modeled as matching it to the empty stem. The skip costs are

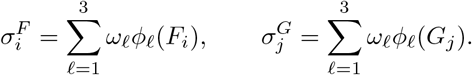

For a terminal coarse alignment represented by *µ* : *D* → *I*_*G*_, the alignment cost is

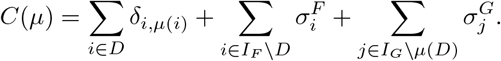

Thus the objective is min_*µ*_ *C*(*µ*), where the minimum is taken over all injective, direction-preserving, and topologically consistent partial maps.

### 4.3 Search states

The algorithm searches over partial alignments in the 5^′^-to-3^′^ order of ℱ. A search node is a triple *u* = (*t, µ, m*), where *t* ∈ {1, …, *f* } is the number of stems of ℱ whose status has already been decided, *µ* is a topologically consistent partial map with domain *D* ⊆ {1, …, *t*}, and

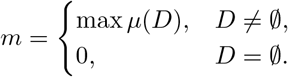

The value *m* is the largest index of a stem in 𝒢 that has been matched so far. By directionality, any unmatched stem *G*_*j*_ with *j < m* is already skipped and cannot be used later.

Equivalently, a node can be represented by two alignment arrays. The ℱ-array has length *f* and stores, for each *i*, either the matched index *µ*(*i*), a skip symbol ⊥, or an undecided symbol ∗. The 𝒢-array is defined analogously. In the implementation these two symbols are encoded by sentinel values; the mathematical description uses ⊥ and ∗.

For a node *u* = (*t, µ, m*), define the fixed cost

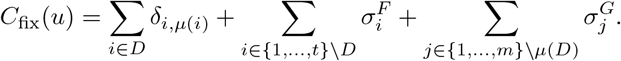

This is the cost of all decisions that can no longer change.

The remaining undecided stems are

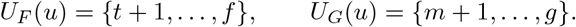

To prioritize the search, we use a lower bound on the cost of completing a partial alignment. For *i* ∈ *U*_*F*_ (*u*), define

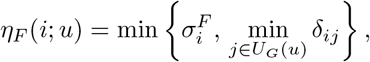

and for *j* ∈ *U*_*G*_(*u*), define

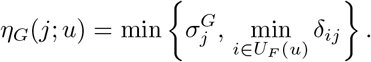

The node key is

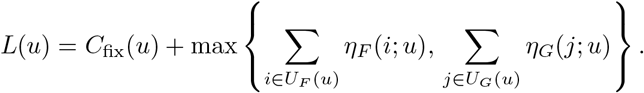

The two sums are not added, since doing so would double-count potential future matches. Their maximum is still a valid lower bound and is sufficient for branch-and-bound pruning.

#### Lemma 4.1.

*For every search node u, L*(*u*) *is a lower bound on the cost of every terminal coarse alignment extending u*.

*Proof*. The term *C*_fix_(*u*) is the exact cost of all matched and skipped stems whose status has already been decided. Consider any terminal extension of *u*. Each remaining stem *F*_*i*_ ∈ *U*_*F*_ (*u*) is either skipped, contributing 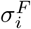, or matched to some remaining stem *G*_*j*_, contributing *δ*_*ij*_. Hence its contribution is at least *η*_*F*_ (*i*; *u*), and the remaining cost is at least 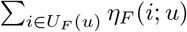. The same argument applied to the remaining stems of 𝒢 shows that the remaining cost is also at least 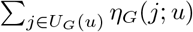. Therefore it is at least the maximum of these two lower bounds. Adding *C*_fix_(*u*) gives *L*(*u*).

### 4.4 Node expansion

Let *u* = (*t, µ, m*) be a non-terminal node, so *t < f* . The next stem to be decided is *F*_*i*_ with *i* = *t* + 1. There is always a skip child (*t* + 1, *µ, m*), whose fixed cost increases by 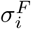.

A match child is generated for a candidate *G*_*j*_, with *j > m*, only if the candidate satisfies all topological constraints induced by the stems already matched in *µ*. Specifically, *F*_*i*_ ↦ *G*_*j*_ is admissible if

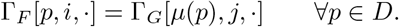

If this condition holds, the child node is

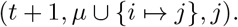

The fixed cost of this child increases by the match cost *δ*_*ij*_ and by the skip costs of any stems of 𝒢 that are passed over:

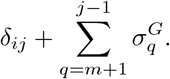

The skipped stems *G*_*m*+1_, …, *G*_*j*−1_ cannot be used later because of the direction-preserving constraint.

Since every pair of matched stems is checked when the later stem in ℱ is added, every terminal node produced by this expansion rule is topologically consistent.

### 4.5 Alignment tree and initialization

The search space is a rooted tree. The root is *u*_0_ = (0, ∅, 0). A root-to-leaf path decides the status of each stem *F*_1_, …, *F*_*f*_ exactly once. When a leaf has *t* = *f*, all remaining unmatched stems *G*_*m*+1_, …, *G*_*g*_ are skipped. Therefore the terminal cost of a leaf *u* = (*f, µ, m*) is

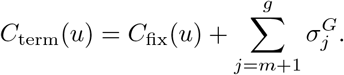

This is equal to the terminal alignment cost *C*(*µ*).

For implementation, we use a shortcut initialization rather than generating leading skip decisions one at a time. The root children are all possible first matched pairs *F*_*i*_ ↦ *G*_*j*_, with 1 ≤ *i* ≤ *f* and 1 ≤ *j* ≤ *g*. In the child corresponding to *F*_*i*_ ↦ *G*_*j*_, stems *F*_1_, …, *F*_*i*−1_ and *G*_1_, …, *G*_*j*−1_ are immediately fixed as skipped. Subsequent expansion then proceeds from *F*_*i*+1_ and considers only stems of 𝒢 with index larger than *j*. Thus the shortcut suppresses paths whose only purpose is to choose the first matched pair after a prefix of skipped stems.

This initialization restricts the implemented search to nonempty partial stem maps: every generated terminal alignment contains at least one matched pair. The all-skip map *µ* = ∅, with cost 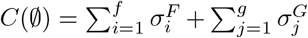, is therefore notgenerated by the shortcut initialization.

### 4.6 Branch-and-bound traversal

The traversal procedure is given in Algorithm 1. The priority queue stores active search nodes ordered by their lower-bound key *L*(*u*). The variable *C*^∗^ stores the best terminal alignment cost found so far, and 𝒮^∗^ stores all terminal alignments attaining this cost. A node is discarded whenever its lower bound exceeds *C*^∗^, since no completion of that node can improve the current best solution. To enumerate all optimal alignments, nodes with lower bound equal to *C*^∗^ are retained until they are either expanded or shown not to lead to another terminal alignment of cost *C*^∗^.

#### Algorithm 1 Best-first branch-and-bound traversal of the alignment tree

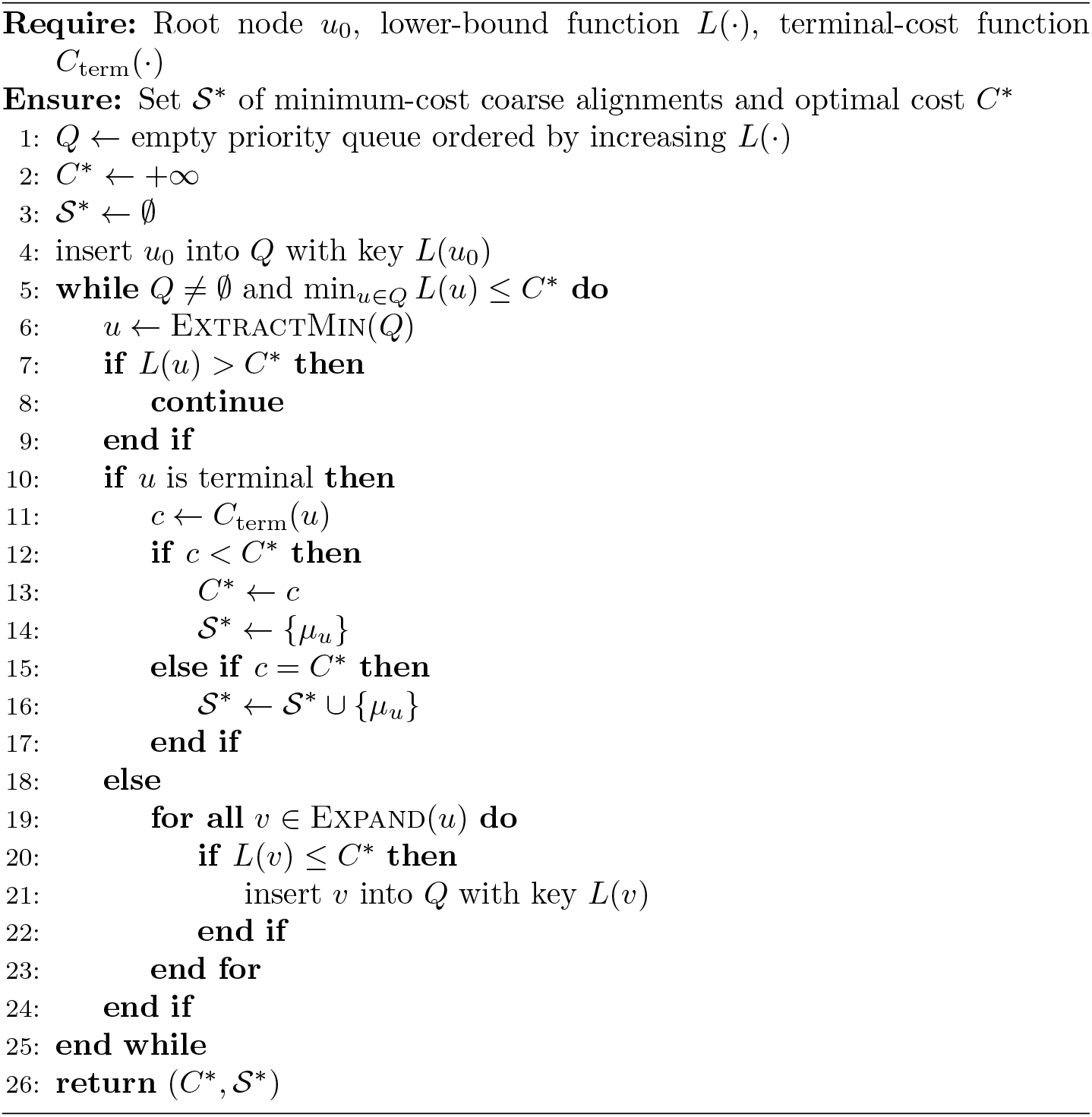

If only one optimal alignment is required, the traversal may stop once a terminal coarse alignment of cost *C*^∗^ has been found and every active node has lower bound at least *C*^∗^. To enumerate all optimal alignments, the search continues to expand nodes with lower bound equal to *C*^∗^, and records every terminal alignment with terminal cost *C*^∗^.

#### Theorem 4.2.

*The branch-and-bound traversal returns a minimum-cost alignment over the set of injective, direction-preserving, and topologically consistent stem maps*.

*Proof*. The expansion rule enumerates exactly the direction-preserving choices for each stem *F*_*i*_: either *F*_*i*_ is skipped, or it is matched to a later unused stem of 𝒢. A match child is generated only when its relation-array entries agree with all previously matched stems, so every terminal node is topologically consistent.

Conversely, any direction-preserving and topologically consistent complete map determines a unique root-to-leaf path in the tree.

By Lemma 4.1, *L*(*u*) is a lower bound on the cost of every terminal coarse alignment extending *u*. Therefore, when a node has *L*(*u*) *> C*^*^, no descendant of that node can improve the best terminal coarse alignment already found. The traversal only prunes such nodes. Hence no alignment with cost smaller than *C*^*^ is discarded. When the queue is empty, or when all remaining nodes have lower bound greater than *C*^*^, no unexplored completion can have cost below *C*^*^. Thus *C*^*^ is the optimal alignment cost.

### Example of topologically constrained expansion

Figure 1 illustrates one expansion step. For a partial alignment with matched-domain *D*, the next stem *F*_*i*_ defines a constraint vector (Γ_*F*_ [*p, i*, ·])_*p*∈*D*_ . For each candidate *G*_*j*_, the algorithm constructs the corresponding vector Γ _*G*_[*µ*(*p*), *j*, ·] _*p*∈*D*_. The candidate *G*_*j*_ is accepted exactly when these two vectors are identical. Otherwise, matching *F*_*i*_ to *G*_*j*_ would change at least one previously established topological relation, and the child is not generated. The skip child for *F*_*i*_ is always generated.

**Figure 1.**
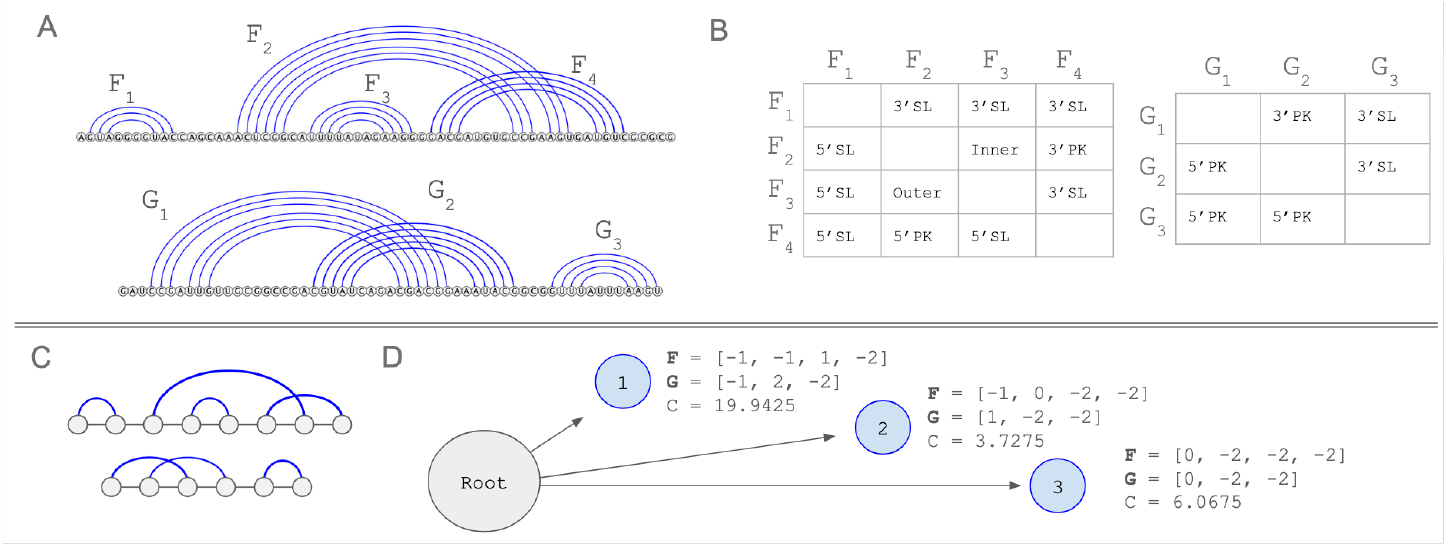
Example of topologically constrained expansion. **A**) Two RNA structures *F* and *G* as arc diagrams (each arc connects two ends of a base pair) with stems labelled. **B**) Relation-array summaries. Rows give the reference stem, and columns give the relation of the column stem with respect to the row stem. Reciprocal entries contain reciprocal relation labels. **C**) Reduced topology representations of *F* and *G* used by the alignment algorithm. **D**) Example search-tree nodes. Each node stores the current partial stem map, skipped stems, undecided stems, and its lower-bound key. A candidate child is generated only if its relation-array constraints match those induced by the previously matched stems.

## 5 Experimental Setup

Table 1 summarizes the Rfam families used in the benchmark and the range of sequence lengths and stem counts after preprocessing.

**Table 1.** Rfam families used in the benchmark and their preprocessing statistics. The first column represent the Rfam family ID; the second column (N. Seq) presents the number of sequences in that family; the third column (Seq. Id.) identifies the sequence similarity in the family; the fourth column (PK) identifies whether the family contains a pseudoknot; the fifth column present the minimum and maximum lengths of sequences in the family; and the last column present the minimum and maximum number of stems in the family as identified in the Rfam structure.

| Rfam family | N. Seq. | Seq. Id. | PK | NT | Stems |
| --- | --- | --- | --- | --- | --- |
| RF00001 | 712 | 0.75 | No | 116-118 | 5-6 |
| RF00003 | 100 | 0.68 | No | 162-164 | 5-5 |
| RF00008 | 85 | 0.71 | No | 40-82 | 3-4 |
| RF00165 | 52 | 0.60 | Yes | 61-63 | 2-2 |
| RF00174 | 434 | 0.73 | Yes | 180-224 | 12-15 |
| RF00381 | 13 | 0.82 | Yes | 56-59 | 2-2 |
| RF00507 | 51 | 0.63 | Yes | 78-81 | 3-4 |

### 5.1 Benchmark instances

For each Rfam family, we selected the first ten sequences and extracted the consensus structure, converting it from WUSS notation into dotbracket and applied the preprocessing procedure described in Section 3. The resulting structures from the same family were not necessarily identical at the stem level: insertions, deletions, and small bulges can change how base pairs are grouped into stems under the chosen bulge compression parameter. We then constructed cross-family alignment instances. For each unordered pair of distinct Rfam families, we formed all 10 × 10 cross-family comparisons by aligning each selected structure from one family with each selected structure from the other. This produced 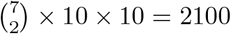 cross-family pairwise alignment instances. We used cross-family rather than within-family pairs because they give a more stringent test of the search procedure: stem features, stem counts, and pairwise topological relations are expected to be less conserved across families.

### 5.2 Measured quantities

For a pair of preprocessed inputs 𝒟_*F*_ = (ℱ, *T*_*F*_, Γ _*F*_, Φ_*F*_), 𝒟_*G*_ = (𝒢, *T*_*G*_, Γ_*G*_, Φ_*G*_), let ℱ = (*F*_1_, …, *F*_*f*_) and 𝒢 = (*G*_1_, …, *G*_*g*_). We use *I*(ℱ, 𝒢) = *fg* as a simple proxy for the number of initial match choices in the shortcut initialization of the alignment tree. This quantity does not determine the full search-tree size, but it captures the first-order dependence on the number of stems in the two inputs.

For each ordered input pair, let 𝒯_ℱ, 𝒢_ be the alignment tree generated by CoSTAR. We recorded the number of generated search nodes and the maximum tree width

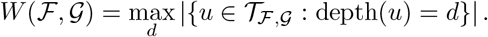

Thus *W* (ℱ, 𝒢) is the largest number of partial alignments present at any search depth. To measure the size of the relation-consistent search space rather than the behavior of a particular branch-and-bound run, we disabled only the cost-based pruning rule *L*(*u*) *> C*^*^ for these measurements. The directionality constraint and the relation-array constraints remained active. Consequently, the reported node counts and tree widths measure the search tree induced by valid topological extensions.

## 6 Results

We evaluated the coarse alignment algorithm on the Rfam benchmark described in Section 5.

### Search-tree growth

Figure 2 summarizes the dependence of running time and tree width on stem-level input size. The left panel plots running time as a function of *I*(ℱ, 𝒢) = *fg*. Instances with larger *I*(ℱ, 𝒢) tend to induce wider search trees and longer running times. This is consistent with the structure of the algorithm: after preprocessing, the dominant cost is not sequence length, but the number of partial stem maps that survive the directionality and topological consistency constraints. The dependence is not solely a function of *f* and *g*, since two pairs with the same stem counts can have different relation arrays and therefore different numbers of admissible extensions. For this panel, the search-driving structure ℱ is set as the structure with more stems.

**Figure 2.**
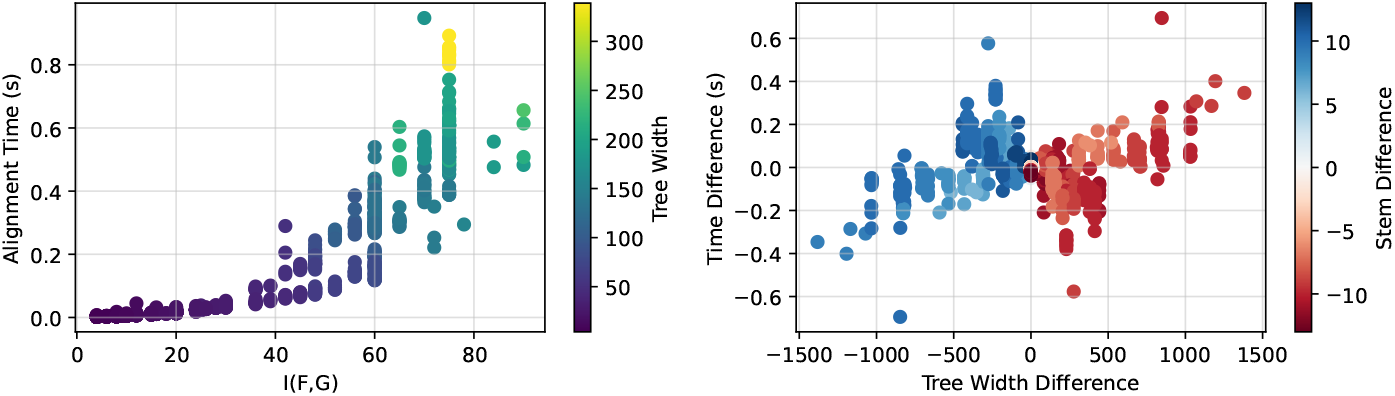
Empirical search behavior on 2100 cross-family Rfam alignment instances. **Left:** Alignment time as a function of the initial stem-pair count *I*(ℱ, 𝒢) = *fg*. Points are colored by maximum tree width *W* (ℱ, 𝒢), computed with cost-based pruning disabled but with directionality and topological constraints enforced. **Right:** Effect of input structure ordering on alignment tree. One structure is set as the search-driving structure (ℱ → 𝒢) and tree width and alignment time are calculated. The reciprocal (𝒢 → ℱ) is also evaluated. For a pair of structures we plot two points showing the change in tree width and alignment time relative to the alternative structure ordering. Color shows which structure has more stems where positive values (blue) indicates the search-driving structure has more stems.

### Effect of input ordering

The alignment problem is symmetric in its final objective, but the search tree is not symmetric in the ordered inputs. The algorithm decides stems of ℱ sequentially and branches over admissible unused stems of 𝒢. Thus, placing the structure with more stems in the search-driving role ℱ increases the depth of the tree but decreases the number of candidate targets considered at each expansion. In the benchmark instances, this ordering reduced maximum tree width. The right panel of Figure 2 compares the alignment tree width and alignment time to calculate ℱ → 𝒢 with the width and time for 𝒢 → ℱ, where ℱ is the structure with more stems. Color indicates the number of stems in ℱ minus the number in 𝒢. When the search-driving structure has more stems, the tree width tends to be lower.

### Preprocessing cost

The non-search preprocessing steps are comparatively inexpensive. For a structure with *n* nucleotides, *p* base pairs, and *M* stems, parsing the structure and computing stem features are linear in the nucleotide-level representation after the base pairs have been ordered. The relation array is computed by comparing all ordered pairs of stems, requiring *O*(*M* ^2^) time and *O*(*M* ^2^) space. Since the relation array Γ_*R*_ and feature list Φ_*R*_ depend only on a single input structure, they can be cached and reused when the same RNA participates in multiple pairwise alignments.

The alignment tree is the main source of computational cost. If relation-array constraints are ignored but directionality is retained, the number of complete direction-preserving partial maps is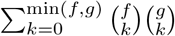 because choosing *k* stems from ℱ and *k* stems from 𝒢 determines a unique order-preserving matching. The generated search tree also contains prefixes of such maps. Relation-array constraints reduce this space in an input-dependent way by rejecting candidate matches that would alter the pairwise topology of already matched stems.

### Adversarial structures

The topological constraints are least effective when many stems have indistinguishable relation profiles. For example, if both inputs contain many repeated stems that are mutually disjoint along the backbone, then most pairwise relations are only 5^′^-disjoint or 3^′^-disjoint. In such cases, relation-array constraints reject few candidate extensions, and the search approaches the directionality-only regime.

As a stress test, we aligned two structures with 11 stems each whose relation patterns were largely repetitive and disjoint. With cost-based pruning disabled, the alignment required approximately six minutes in our implementation, generated about 10^6^ search nodes, and reached a maximum tree width of approximately 2.8 × 10^5^. This instance illustrates the data-dependent nature of the search: the number of stems determines the broad scale of the search tree, but the pairwise topology of those stems determines how much pruning is obtained from the relation-array constraints.

### 6.1 Qualitative comparison with LaRA

We first compare CoSTAR with LaRA, the closest available method for aligning RNA sequences with pseudoknotted secondary structures. LaRA optimizes a nucleotide-level sequence–structure alignment, whereas CoSTAR optimizes a partial map between stems. Thus, the two methods do not return the same type of object. The examples below show that, even when LaRA is run in a structure-prioritized setting, its nucleotide-level alignment can fail to recover the stem correspondences required for coarse structural comparison.

#### Synthetic pseudoknotted pair

We constructed a synthetic pair in which the two RNAs share a pseudoknotted structural core but differ in flanking and additional stems. This design tests whether an aligner can ignore non-corresponding structural elements and align the shared pseudoknotted block. Listing 1 shows the LaRA alignment. The shared paired regions are not aligned as coherent stem blocks: gaps are inserted within paired regions, and the pseudoknotted arms are fragmented across the alignment. As a result, the output does not provide a usable stem-to-stem correspondence for this example.

Listing 1: LaRA nucleotide-level alignment of the synthetic pseudoknotted pair.

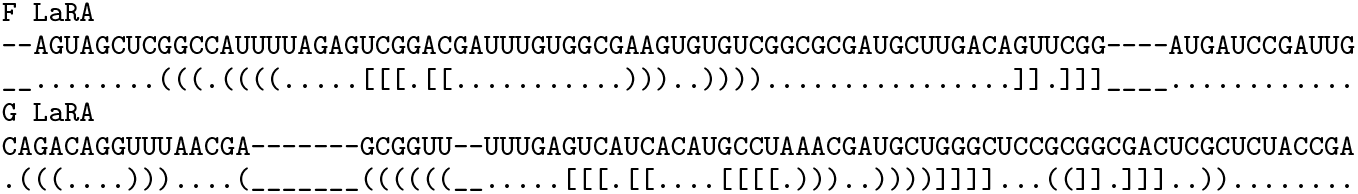

Listing 2 shows the nucleotide-level projection obtained from the CoSTAR stem map on the same input pair. Vertical bars mark the boundaries induced by the coarse alignment. The shared pseudoknotted arms are aligned within corresponding blocks, while additional stems and unmatched regions are separated by gaps or distinct blocks. This illustrates the intended use of CoSTAR: the structural correspondence is determined at the stem level before nucleotide columns are introduced.

Listing 2: CoSTAR nucleotide-level projection of the synthetic pseudoknotted pair. Vertical bars mark blocks induced by the coarse stem map.

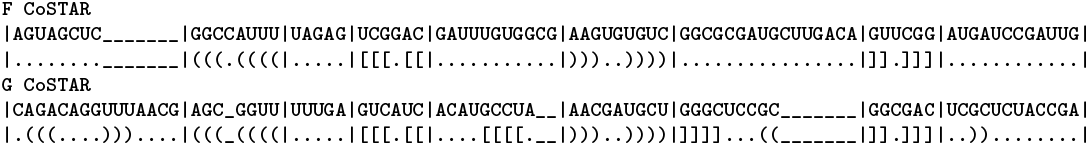

#### Stem-deletion perturbation

We next tested whether the methods can align a structure to a perturbed copy with a deleted stem. We selected a structure from Rfam family RF00507 and created a knockout by removing the first stem together with external unstructured nucleotides. The desired coarse alignment should skip the deleted 5^′^ stem and align the remaining stems after a leading gap.

Listing 3 shows the alignment produced by LaRA. LaRA introduces a large 5^′^ gap, but the gap is placed within the remaining paired region rather than before the corresponding structural block. Consequently, the deleted stem is not represented as a clean skipped structural element, and the alignment does not recover the intended stem-level offset.

Listing 3: LaRA nucleotide-level alignment of the RF00507 stem-deletion perturbation.

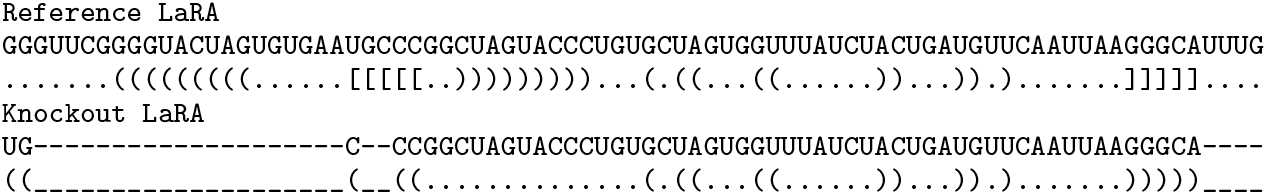

CoSTAR instead represents the deletion as a skipped leading block and aligns the remaining stems directly, as shown in Listing 4. This is the behavior required for the coarse alignment task: the perturbation is localized to the deleted stem, while the rest of the stem map remains interpretable.

Listing 4: CoSTAR nucleotide-level projection of the RF00507 stem-deletion perturbation. The deleted leading stem is represented as a skipped block.

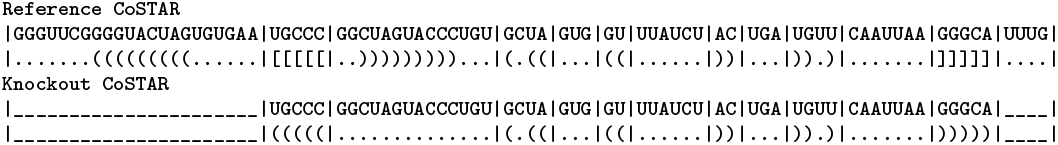

Together, these examples demonstrate the distinction between the two objectives. LaRA can align pseudoknotted RNA at nucleotide resolution, but the resulting alignment need not preserve the stem-level correspondences needed for coarse structural comparison. CoSTAR directly optimizes that stem-level correspondence and can then project the resulting map to nucleotide resolution.

## 7 Discussion and Future Work

We presented CoSTAR, a coarse alignment algorithm for RNA secondary structures that may contain pseudoknots. CoSTAR represents each input by its ordered stems, pairwise stem relations, and nucleotide-derived stem features, and returns a partial injective map between stems. The method therefore lies between topology-only representations, such as RNA abstract shapes and RNA-as-Graph [13, 11], and nucleotide-level sequence–structure alignment methods such as LaRA 2 [28]. Its main computational advantage is that the dominant input size is the number of stems, not the number of nucleotides: stem features are computed once, relation arrays require *O*(*M* ^2^) time and space for *M* stems, and these precomputed objects can be cached across alignments.

The search over partial stem maps remains the computational bottleneck and can grow exponentially in the number of stems in the worst case. The empirical results show, however, that directionality and relation-array constraints reduce the explored search space on the benchmark instances. Because the search is asymmetric in its ordered inputs, using the structure with more stems as the search-driving input ℱ typically reduces tree width, as observed in the reciprocal-orientation experiments. The least favorable cases are structures with many repeated, mutually disjoint stems, where most relations are 5^′^-disjoint or 3^′^-disjoint and topological constraints reject few candidates.

Several modeling choices can be refined. The current lower bound is admissible but relaxed: it allows each remaining stem to choose its cheapest match or skip cost without enforcing future injectivity or topological consistency. Tighter admissible bounds, such as a minimum-cost bipartite matching over the remaining stems with skip vertices, could reduce the number of expanded nodes. The cost function is also intentionally simple, using weighted distances between stem features and corresponding skip costs. Since RNA similarity is task-dependent [3], future scoring models could incorporate sequence composition, covariation, learned feature weights, or stronger penalties for disrupting pseudoknotted topology.

CoSTAR can also be used as an anchoring step for nucleotide-level alignment. If the coarse map matches *F*_*i*_ to *G*_*j*_, then the nucleotide positions in the arms of *F*_*i*_ can be constrained or encouraged to align to the corresponding arms of *G*_*j*_, and intervals between matched stems can be solved as smaller sequence–structure alignment subproblems. We provide a simple nucleotide-projection method, though future work should evaluate this projection on curated homologous RNAs, viral pseudoknot benchmarks, and comparisons against nucleotide-level aligners and topology-based distance methods. Additional extensions include studying the effect of the bulge-compression parameter *β*, which trades off fewer stems and faster search against the risk of merging stems that should remain distinct.

## A Examples of Nucleotide-Level Alignment Behavior

LaRA 2 was used as a point of comparison because it supports alignment of RNA sequences with pseudoknotted secondary structures [28]. However, LaRA solves a nucleotide-level sequence–structure alignment problem, whereas our method computes a stem-level structural map. The two outputs are therefore not identical objects. The examples below illustrate cases in which a nucleotide-level alignment can be difficult to interpret when the intended comparison is a coarse stem correspondence.

Let *S*^*F*^, *R*^*F*^ and *S*^*G*^, *R*^*G*^ denote the two input sequence–structure pairs. A nucleotide-level aligner returns aligned strings 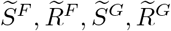 where gap symbols are inserted into the sequence and structure strings. A base pair (*i, j*) ∈ *R*^*F*^ is preserved by the alignment if the columns containing 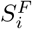 and 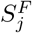 are aligned to two non-gap nucleotides 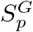 and 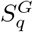 such that (*p, q*) ∈ *R*^*G*^. In the examples below, we use this criterion only as a qualitative diagnostic: the purpose is not to benchmark LaRA, but to show why a stem-level output can be more appropriate for some structural comparison tasks.

LaRA has three relevant parameters in these examples. The parameter `balance` controls the relative contribution of sequence and structure. We set `balance=INF` to prioritize structural agreement. The parameters `gapOpen` and `gapExtend` define the affine cost for opening and extending gaps, respectively.

### Default gap parameters

The first run used the two structures shown in Figure 1, with `balance=INF` and the default LaRA gap parameters. Lines labelled *F* and *G* give the aligned sequence and structure strings for the two inputs.

Listing 5: LaRA output with`balance=INF` and default gap parameters.

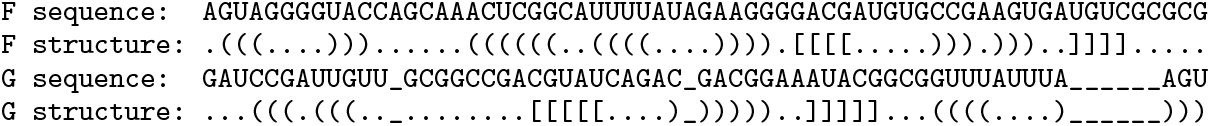

In this output, gaps are inserted inside structural regions of *G*. For example, an insertion occurs within a bracketed segment of the *G* structure, splitting a stem arm in the aligned representation. A second long insertion appears near the end of the structure string. As a result, the displayed alignment does not preserve the intended correspondence between the stem arms and pseudoknotted arms visible in Figure 1. This is a nucleotide-level behavior relative to the stem-level objective considered in this paper: the alignment is syntactically valid, but it does not provide a useful mapping between stems.

### More permissive gap parameters

We repeated the same comparison with lower gap costs, using `gapOpen=3` and `gapExtend=1.` This makes long gaps less costly and should, in principle, allow the nucleotide-level alignment to place larger insertions outside conserved structural blocks.

Listing 6: LaRA output with `balance=INF`, `gapOpen=3`, and `gapExtend=1.`

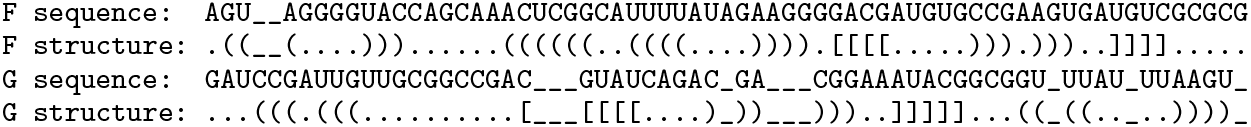

Lowering the gap penalties does not remove this behavior in the example. Instead, gaps are still placed inside bracket runs, and the resulting alignment continues to split regions that should be treated as coherent stems. This behavior reflects the difference between optimizing a nucleotide-level alignment objective and directly optimizing a map between stems.

### Projection of the coarse alignment

For comparison, Listing 7 shows a nucleotide-level projection of the coarse alignment produced by our method. The method first computes a partial map between stems. Matched stems are then projected to nucleotide resolution by aligning corresponding stem arms as blocks. The vertical bars mark the boundaries of coarse stem blocks.

Listing 7: Nucleotide-level projection of the stem-level alignment. Vertical bars mark coarse stem boundaries.

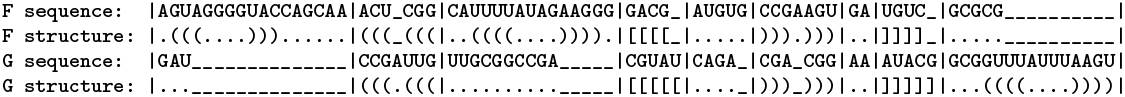

The projection is not intended to replace a full nucleotide-level sequence–structure aligner. Rather, it shows the type of information produced by the coarse algorithm: corresponding stems are identified first, and nucleotide-level columns are introduced only after the stem map has been fixed. This avoids placing gaps inside a stem arm when the downstream task is to compare stems as coherent units.

In other LaRA runs, we also observed a 5^′^-anchoring behavior, where an early nucleotide was aligned before a leading gap even though introducing the gap would better align the downstream stems. Such cases further motivate the use of stem-level alignment as a preprocessing or anchoring step when the primary biological question concerns structural correspondence rather than nucleotide identity.

## B Rfam Samples

In our results we evaluated our method on Rfam RNA structures, specifically the first 10 of each family. Table 2 contains the accession numbers for the used samples.

**Table 2.** Rfam family sample accession numbers.

| **RF00001** | **RF00003** | **RF00008** | **RF00165** | **RF00174** | **RF00381** | **RF00507** |
| --- | --- | --- | --- | --- | --- | --- |
| X01556.13-118 | X06810.1261-421 | D00685.11-46 | NC_022103.127934-27996 | AF010496.139869-39653 | BC056833.1241-299 | NC_006577.213601-13681 |
| X55260.13-119 | X14417.1177-340 | AJ247116.1133-53 | NC_009657.127966-28028 | AF010496.1105318-105540 | AF242520.1225-280 | NC_003045.113342-13422 |
| M16174.13-119 | Z11883.13441-3603 | Y14700.1133-53 | NC_028814.127384-27446 | AF010496.1116971-117193 | AF258463.1230-288 | NC_006213.113341-13421 |
| X55267.13-119 | X14419.1178-338 | AJ247123.1132-52 | NC_038294.129868-29928 | AF193754.14343-4143 | D10706.1256-314 | NC_026011.113485-13565 |
| M16172.13-119 | X14416.1171-331 | AJ247121.1133-53 | NC_019843.329875-29935 | AF193754.124966-24789 | AC004152.116641-16699 | NC_004718.313399-13476 |
| AF001265.16033-6149 | X14415.1175-335 | AJ247122.1132-52 | NC_003436.127820-27882 | AF263012.19229-9016 | AF291576.1215-270 | NC_022103.112533-12607 |
| X05057.13-119 | X14414.1152-312 | AJ247113.1134-53 | NC_038861.128363-28425 | AF306632.1382-182 | D11373.1473-530 | NC_002306.312410-12484 |
| X15126.13-120 | Z11883.12839-2998 | AJ536620.11-40 | NC_028806.127893-27955 | AJ000758.11191-1369 | BC054690.1204-259 | NC_038861.112339-12413 |
| Z50737.13-119 | Z11883.11496-1656 | AJ536617.11-40 | NC_002306.329138-29200 | AJ000758.11489-1667 | AB017117.1228-286 | NC_028806.112325-12399 |
| X55261.13-119 | Z11883.1361-521 | Y12833.1339-285 | NC_032107.128249-28311 | AJ278144.122351-22164 | AF196332.169-127 | NC_017083.113520-13600 |

